# AtlasAgent: Vision language model and Agent-guided Framework for Evaluation of Atlas-scale Single-cell Integration

**DOI:** 10.1101/2025.07.15.663271

**Authors:** Danqing Yin, Zhongmin Zhang, Xinci Liu, Ke Ni, Huidong Su, Nicolas Lin Li, Hongyu Dong, Qiuchen Zhao, Xinyi Lin, Luyi Tian, Ye Meng, Joshua W K Ho

## Abstract

As single-cell omics transitions into the era of AI-virtual cells (AIVC), where large-scale single-cell data integration becomes prevalent, the computational demands of integration evaluation emerge as critical scalability bottlenecks. Traditional integration evaluation pipelines, requiring metrics like k-nearest-neighbor batch effect test (kBET) and Local Inverse Simpson’s Index (iLISI) employed by state-of-the-art scIB method, often demand large computational resources and long runtimes, making them infeasible for large scale integration studies. Herein, we present AtlasAgent, the first vision-language model (VLM)-powered and AI agent framework to accelerate atlas-scale integration evaluation at unprecedented speed and scale. We systematically evaluate batch correction quality, biological signal preservation and overcorrection risks using chain-of-thought reasoning in conjunction with few-shot and zero-shot prompting strategies. AtlasAgent completes evaluation within 32 seconds, in contrast to scIB’s runtime of 5.55 hours in GPU, while identifying the scIB-determint best integration methods within the top-3 in 88.3% of the time, lowering evaluation time from hours to seconds while preserving alignment with domain expert reasoning. AtlasAgent pioneers the use of VLMs to realize scalable and rapid integration evaluation at atlas scale.

## Introduction

The integration of single-cell omics datasets across batches, conditions, or technologies is a foundational task in single-cell bioinformatics [1]. Data integration refers to the processing of cells from different sources that can be placed in the same embedding, in order to measure distances between them, allowing the cell type annotation, and additional downstream joint analyses. As single-cell datasets grow in size and diversity in the AIVC era, building AIVC requires reliable batch effect correction and rigorous evaluation mechanisms for multi-modal single-cell data integration [2]. Single-cell integration methods aim to remove technical batch effects while preserving biological variation, therefore a critical measure of integration quality is based on how well the integration algorithm balances the tradeoff between the two factors [6]. While integration algorithms have proliferated rapidly [3–5], the field continues to face a critical bottleneck: the quantitative evaluation of integration quality remains computationally demanding [6], manually intensive, and conceptually dataset-specific based on divergent metrics [3,6].

Numerous metrics have been developed to assess batch correction and biological conservation. The scIB framework [6] has formalized a benchmark suite comprising 14 metrics, including batch mixing metrics (kBET, graph iLISI, graph connectivity, PC regression) and biological conservation metrics (NMI, ARI, ASW (cell types), graph cLISI, isolated label F1, isolated label silhouette, HVG overlap, cell cycle variance, trajectory conservation, and cell type ASW). While scIB provides a structured foundation for integration assessment, it inherits limitations such as scalability and usability shared across earlier frameworks [6]—notably, the high computational overhead, reliance on ground truth labels, and lack of interpretability for end users.

Furthermore, mainstream evaluation tools rely on statistical testing or label-dependent performance metrics, which may obscure biologically relevant structure or over-penalize integration methods in complex tissues with gradual transitions [5,7]. Specifically, statistics-based frameworks including scIB have been shown to be restrained, focusing only on dataset-specific measures with the risk of neglecting dataset-agnostic (e.g., visual cues) features in developmental or trajector-rich single-cell atlases [7, 8].

Recent advances in large language models (LLMs) and autonomous AI agents have opened new directions for automating complex analytical tasks in computational biology [9–11]. While the use of LLMs and agents in single-cell analysis workflow automation has gained traction [12–16], their application in single-cell atlas-scale integration remains nascent. Emerging studies demonstrate LLMs’ capacity to democratize single-cell analysis through natural language-guided automation—generating code, executing pipelines, and performing exploratory analysis [11]—while also enabling prior-guided batch correction [17], though their application to integration evaluation remains nascent. Large-scale integration tasks are challenging to perform evaluation, it is time to move beyond traditional computation-heavy metrics-based methods to a vision-based approach.

To systematically address these challenges and unmet needs in the current frameworks, we introduce AtlasAgent+, the first task-autonomous, agent-driven evaluation framework for single-cell data integration. Rather than relying on conventional numeric metrics, AtlasAgent+ employs a vision-language paradigm wherein 2-D scatterplot such as UMAP visualizations are treated as primary evaluative substrates [9,10], which is commonly used in evaluation of data integration. The framework is embedded in an agent, integrating modules for visual reasoning [9,10], retrieval-augmented memory, and self-consistency voting, enabling assessments across three principal axes: batch mixing, biological conservation, and overcorrection risk.

Each agent in the AtlasAgent+ framework has been assigned a distinct functional role with dedicated and fine-grained prompts—(1) CriteriaExtractorAgent for visual-metric alignment and visual criterion extraction [9], and (2) VisionEvaluatorAgent for visual evaluation of the UMAP visualization from integration results. Together, they contribute to a collaborative reasoning loop modeled after expert analytical workflows [15,16]. This structure eliminates the need for manual metric selection, complex statistical computation, and labeled ground truths, allowing evaluation within seconds even on large-scale datasets.

AtlasAgent+ further supports method selection, adapting dynamically to dataset-specific properties and integrating contextual memory from prior evaluations. Validated across diverse integration scenarios—including cross-tissue [6], cross-technology platform [7], cross-species [5], and trajectory-resolved benchmarks [8]—AtlasAgent+ demonstrates alignment with expert judgments while dramatically reducing computational burden and enabling autonomy and scalability.

By reframing integration evaluation as a goal-driven, agent-based reasoning task, AtlasAgent+ represents a next generation tool addressing atlas-level single-cell data integration evaluation. It offers a scalable, interpretable, and robust solution to long-standing bottlenecks in scIB e.g. scalability and usability.

## Method

### Benchmark Dataset Selection and Integration Task Design

To construct a diverse evaluation framework for atlas-scale integration, we curated a representative collection of single-cell datasets’ integration UMAP plots spanning a variety of tissues, species, and experimental platforms. A total of 7 integration tasks were performed for atlas-scale scenarios characterized by complex batch effects, varying cell type compositions, and increasing cell counts. The UMAP plots come from two primary sources—peer-reviewed publications and curated online repositories [19, 20].

First, we begin by harvesting raw count matrices and associated metadata from curated online repositories, raw datasets then undergo quality control and normalization, followed by data integration in six state-of-the-art integration algorithms (Harmony, Scanorama, BBKNN, SAUCIE, scVI and scANVI). The outputs of these methods served as the basis for both scIB metric-based and agent-based evaluation. To establish quantitative ground truth, we employed the scIB benchmarking framework, which evaluates integration results across 14 standardized metrics. These include batch removal metrics (kBET, graph iLISI, graph connectivity, PC regression) and biological conservation metrics (NMI, ARI, ASW (cell type), graph cLISI, isolated label F1, isolated label silhouette, HVG overlap, cell cycle variance, trajectory conservation, and cell type ASW). All integration outputs were benchmarked using scIB under consistent preprocessing settings. Scores were recorded per metric for each method-dataset pair.

On the other hand, UMAP plots of integration tasks were sourced and extracted from public literature..

We compiled 84 composite figures. Each figure contains an “Unintegrated” reference panel and between seven and varying number of integration methods. The figures fall into diverse categories—PBMC, pancreas, lung, retina, whole-embryo, cell-line mixtures and haematopoietic progenitors —— ensuring diversity in batch size (2 – 8 studies) and cell-type complexity (2 – 15 annotated types).

For every panel we extracted the **overall** column winner(s) of integration method from the original scIB evaluation report or other evaluation method of integration quality. A meta-table listing task descriptor, number of batches, number of cell types and statistically evaluated gold winner(s) is provided in Supplementary Data 1.

**Figure 1.**
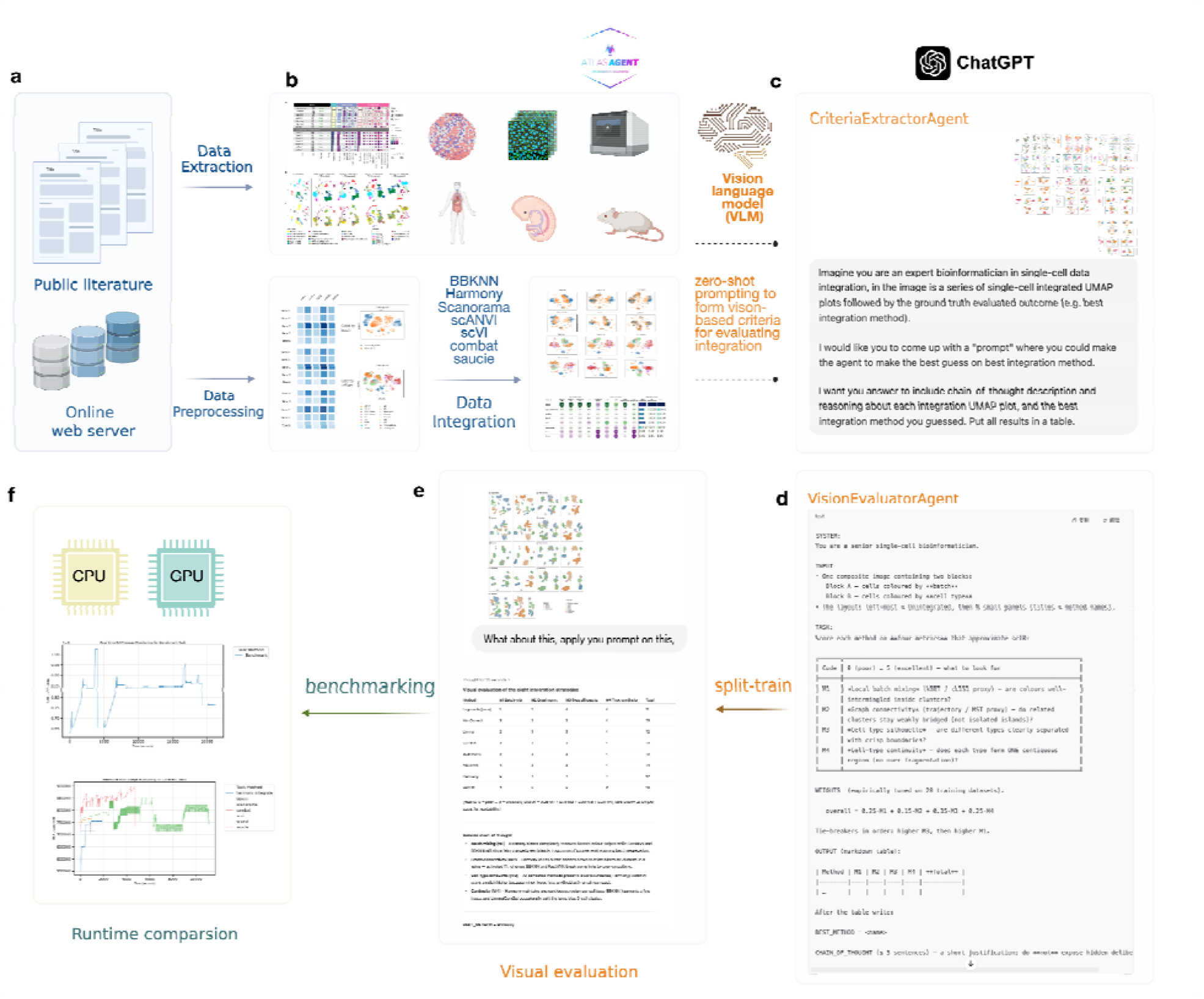
AtlasAgent+: A vision-language, LLM-driven framework for automated single-cell integration evaluation. Overview of the AtlasAgent+ framework workflow and benchmarking results. Starting from a curated literature dataset (**a**), AtlasAgent+ employs vision-language reasoning on UMAP visualizations and integration metrics extracted from training panels (**b**). Tasks are evaluated by two sub agents against ground truths, with AtlasAgent+ generating automated predictions via integrated OpenAI APIs (**c; d**). Benchmarking comparison among various GPT model variants (**e**) in assessing integration quality.

### 2. Generation of Vision-Based Evaluation Standards via LLM-Agent Interaction

To develop a vision-language-based alternative to statistical evaluation like scIB, we designed a structured prompting protocol termed the CriteriaExtractorAgent. Using the GPT model as the underlying language model, we exposed the agent to two-dimensional UMAP visualizations derived from integrated outputs. Prompts were programmatically constructed to elicit visual pattern recognition aligned with batch mixing quality, biological structure retention, and overcorrection patterns as considered by traditional method scIB. Each agent interaction consisted of a multi-turn reasoning loop using few-shot examples to learn metric-visual pattern equivalence.

We employed a series of latest vision language models such as GPT-4o, GPT-o3, GPT-4o-mini and GPT-4-turbo with API for automated vision-based batch evaluation to compare the computational accuracy as well as efficiency among them.

#### Few-shot prompting

The visual rubric was established through an **in-context few-shot procedure**: 7 representative composites, each accompanied by its scIB overall winner, were chosen based on domain experts for balancing datasets that cover common scenarios in single-cell biology. The 4 visual criteria (M1–M4) and a dedicated weight for scoring each criteria were developed. After this calibration step the rubric was applied **zero-shot** to the remaining composites, i.e. the model received no additional labels. During every evaluation the agent produced a brief **chain-of-thought explanation** that justified the numeric scores in natural language; this explicit reasoning improved internal consistency and offered readers an interpretable audit trail.

#### Calibration phase

At the very start of the dialogue we supplied **7 fully-labelled UMAP visualization panels** (each with its scIB overall winner explicitly shown). These 7 examples were referenced repeatedly while we wrote and refined the prompts. Within the LLM this constitutes **in-context few-shot learning**: the model inferred how high or low to rate each visual cue (M1–M4) and discovered that two particular weight vectors (equal-weight v1 and scIB-aligned v2) reproduced the exemplar winners most reliably.

#### Zero-shot generalisation phase

Once the two prompts were frozen, the model was asked to score a further remaining composites that it had never ‘seen’ with labels, this becomes our **VisionEvaluatorAgent**. We define *k* to be 3 to allow the agent to pick top-*k* best integration method(s) as compared to gold-standard ranked integration methods by scIB and other statistical tools. For those panels the model operated in a zero-shot regime: it applied the rubric and the previously chosen weights without being given the answer in advance. All accuracy statistics (winner agreement) were collected for summarization to compare agent performance versus traditional evaluation tools.

#### Chain-of-thought reasoning

For every panel the VisionEvaluationAgent produced a short narrative—typically one to three sentences—explaining why the top-ranked method was preferred (“batch colours are fully intermingled while cell-type islands remain single-colour” etc.). This explicit explanation is classic chain-of-thought prompting; it forces the model to articulate intermediate observations before finalising the verdict. Empirically, asking for such reasoning reduced hallucinated winners and improved inter-run stability..

#### Logical architecture of AtlasAgent used in this study

AtlasAgent was conceived as a dual-agent design:

1. **CriteriaExtractorAgent** is responsible for perception. It inspects a composite UMAP figure and verbalises salient visual cues or criteria for integration evaluation.
2. **VisionEvaluatorAgent** is responsible for visual judgement of UMAP visualization panels. It transforms the CriteriaExtractorAgent description into numerical scores using a fixed visual rubric and returns a ranked table together with a short justification.

In the present simulation both roles were executed sequentially by querying the GPT model. The model first generated an internal description (**CriteriaExtractor**Agent-like) and immediately used that description to compute rubric scores (VisionEvaluationAgent-like). An additional ChatGPT API wrapped in Python code was invoked for batch processing of UMAP panel images.

### Runtime Profiling of Metric-Based Evaluation Pipelines

A Scalability comparison

To quantify the computational cost of traditional evaluation strategies, we benchmarked the execution time of scIB metrics across varying data scales. We evaluated runtime across dimensions of total cell count, batch number, and method complexity. Results were used to construct efficiency scaling profiles for comparison with the instantaneous inference time of EvaluatorAgent.

#### Software, Infrastructure, and Reproducibility

All experiments were conducted using Python 3.10 backends. LLM inference was powered by ChatGPT via OpenAI API. Experiments were executed in two high-performance computing GPU-powered servers (NVIDIA 1XA100 GPUs with 1TB RAM and NVIDIA 4XA100GPUs with 600GB RAM).

Agent prompts, evaluation logs, and datasets are publicly available at https://github.com/hiyin/AtlasAgent.

## Results

### Integration quality varies across methods and atlas-level scenarios

We take 7 UMAP visualizations as calibration set to derive the visual criteria, i.e. VisionEvaluatorAgent. The 7 figures correspond to atlas-scale integration tasks encompassing a range of biological contexts and technical complexities, including cross-cohort COVID-19 datasets, cross-tissue developmental atlases, and aging cell landscapes. Each task was integrated using 6 representative methods spanning from commonly favored methods to deep learning based methods: Harmony, SAUCIE, BBKNN, Scanorama, scANVI and scVI.

### Accuracy and assessment

The test set is composed of the remaining 77 panels, outside of the 7 calibration UMAP composite. The **VisionEvaluatorAgent** backed by ChatGPT-4o matched the gold-standard integration method top-3 winners in 88.3 % of the panels, whereas ChatGPT-o3 matched in 82.5% panels. ChatGPT-4o-mini produced 60 correct answers at 77.9 %, while GPT-3.5-Turbo produced 56 correct answers at 72.7 %.

**Figure 2:**
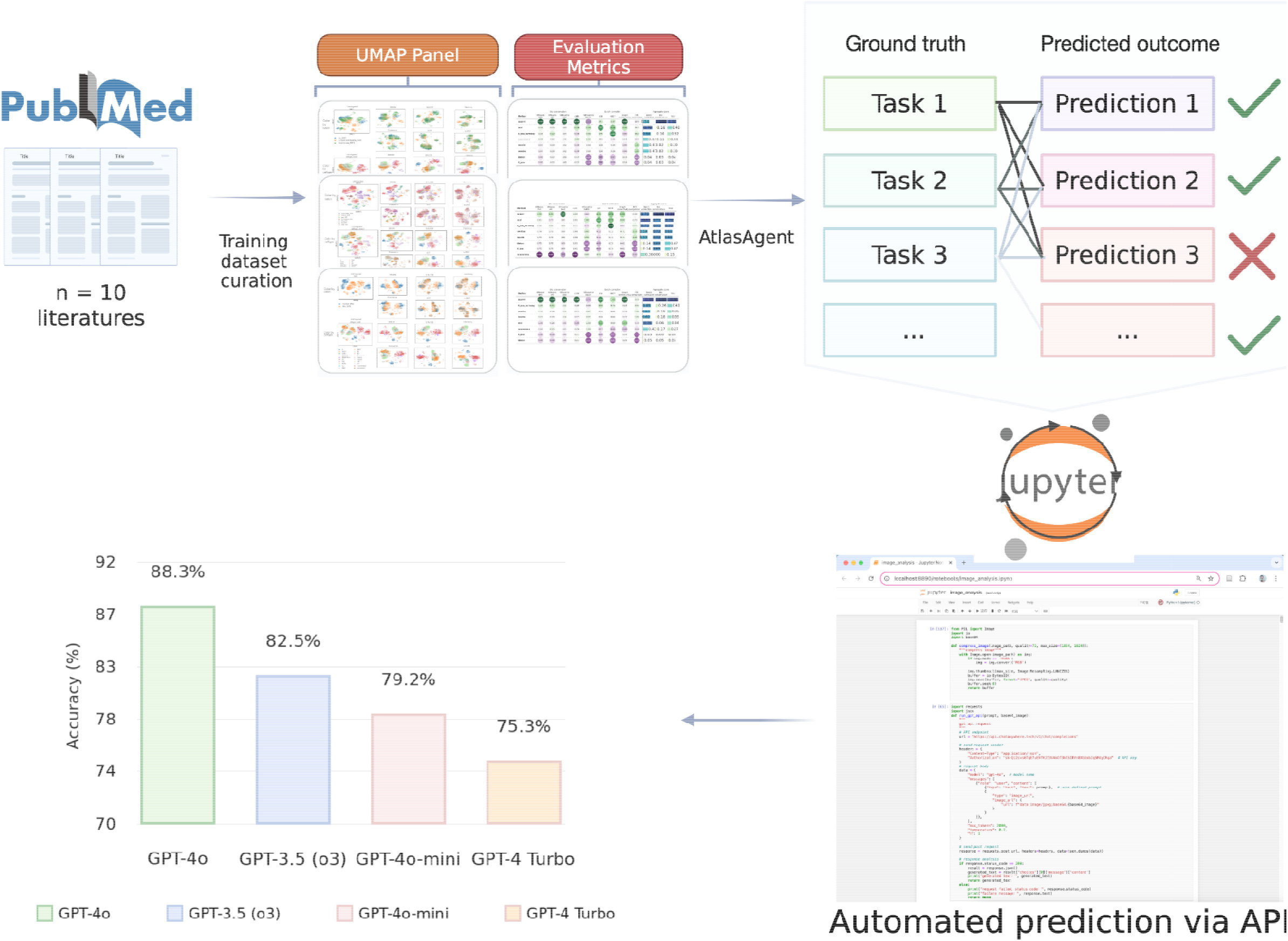
AtlasAgent+: A vision-language, LLM-driven framework for automated single-cell integration evaluation Overview of the AtlasAgent+ framework workflow and benchmarking results. Starting from a curated literature dataset, AtlasAgent+ employs vision-language reasoning on UMAP visualizations and integration metrics extracted from training panels (top-center). Tasks are evaluated against ground truths, with AtlasAgent+ generating automated predictions via Jupyter-integrated APIs. Benchmarking comparison among various GPT model variants demonstrates GPT-4o’s superior accuracy in assessing integration quality.

### Derivation of the visual rubric (M1 – M4) and weight sets

To identify a minimal, human-interpretable proxy for the 14 numeric criteria used by scIB, we performed an inductive analysis on the **7 calibration composites** supplied at the outset of this study. ChatGPT-o3, prompted to derive a series of visual cues for evaluating the UMAP figure.

#### VisionExtractorAgent prompt

Imagine you are an expert bioinformatician in single-cell data integration, in the image is a series of single-cell integrated UMAP plots followed by the ground truth evaluated outcome (e.g. best integration method). I would like you to come up with a “prompt” where you could make the agent to make the best guess on best integration method. I want you answer to include chain-of-thought description and reasoning about each integration UMAP plot, and the best integration method you guessed. Put all results in a table.

These observations were recast in positive form to yield 4 visual cues as summarized by GPT model:

Table 1 illustrates each cue with side-by-side “good” and “bad” exemplars extracted from the calibration panels.

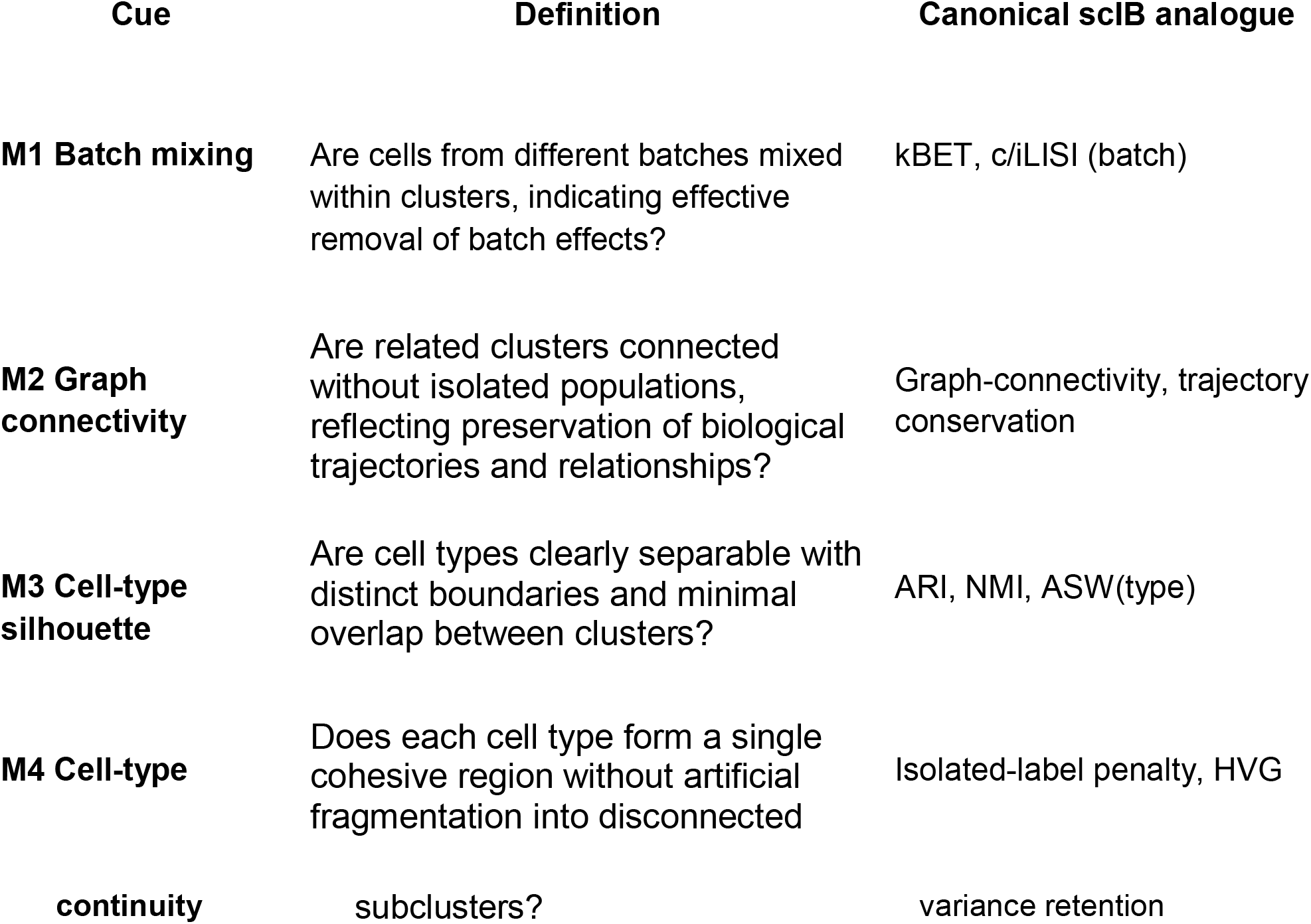
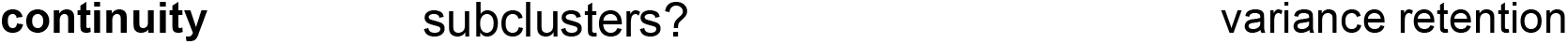

A pilot scoring round on the same 7 panels showed that **four scIB-aligned cues were sufficient to identify the scIB-equivalent winner in every case**. The 7 calibration UMAP panels were scored inside the chat session, and two practical weight schemes were selected by manual trial-and-error to maximise concordance with the printed scIB tables:

- **Prompt weights**

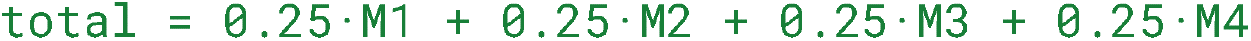

No external optimisation code or additional human graders were used; all agent-based evaluation decisions occurred within the chat session.

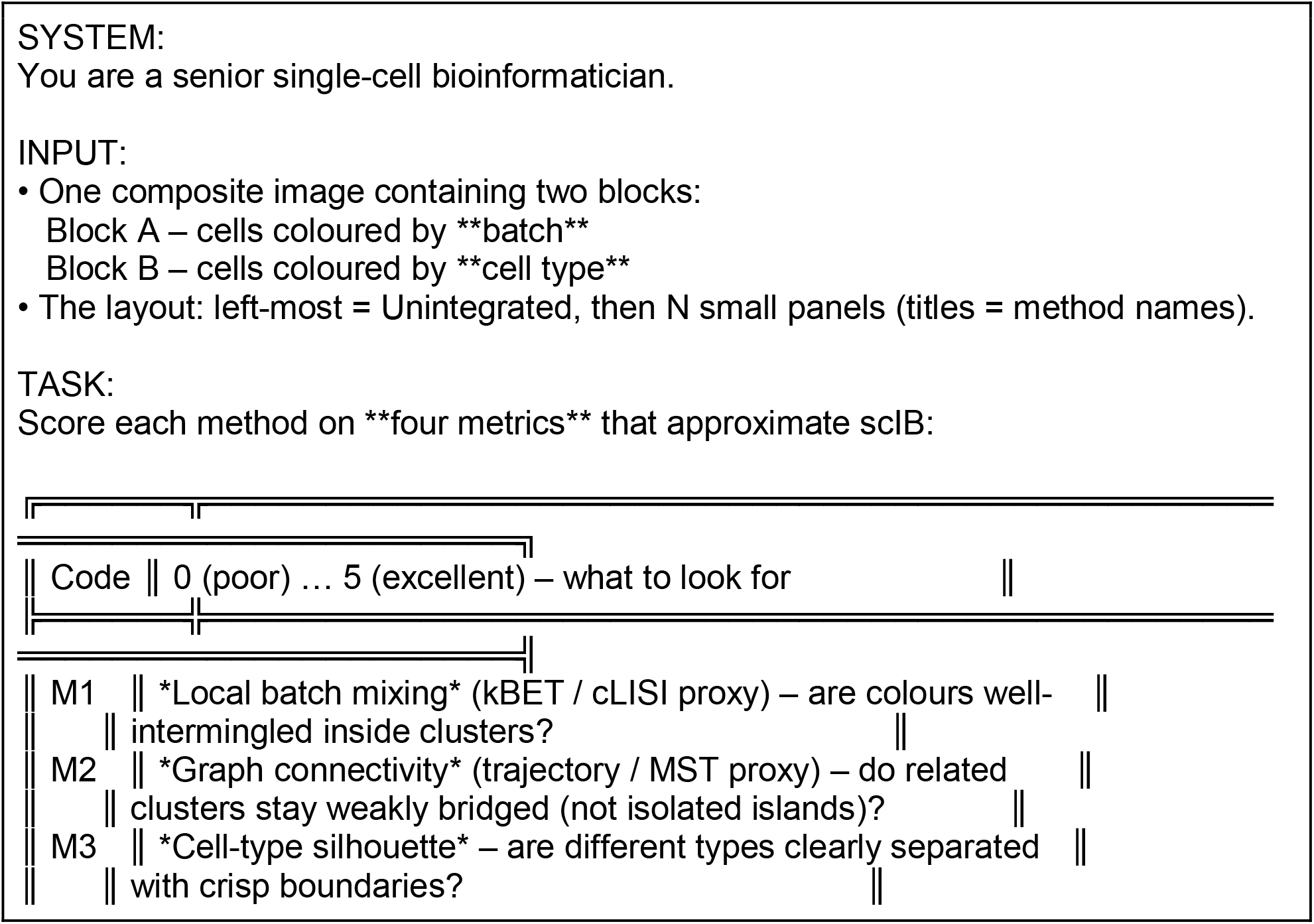
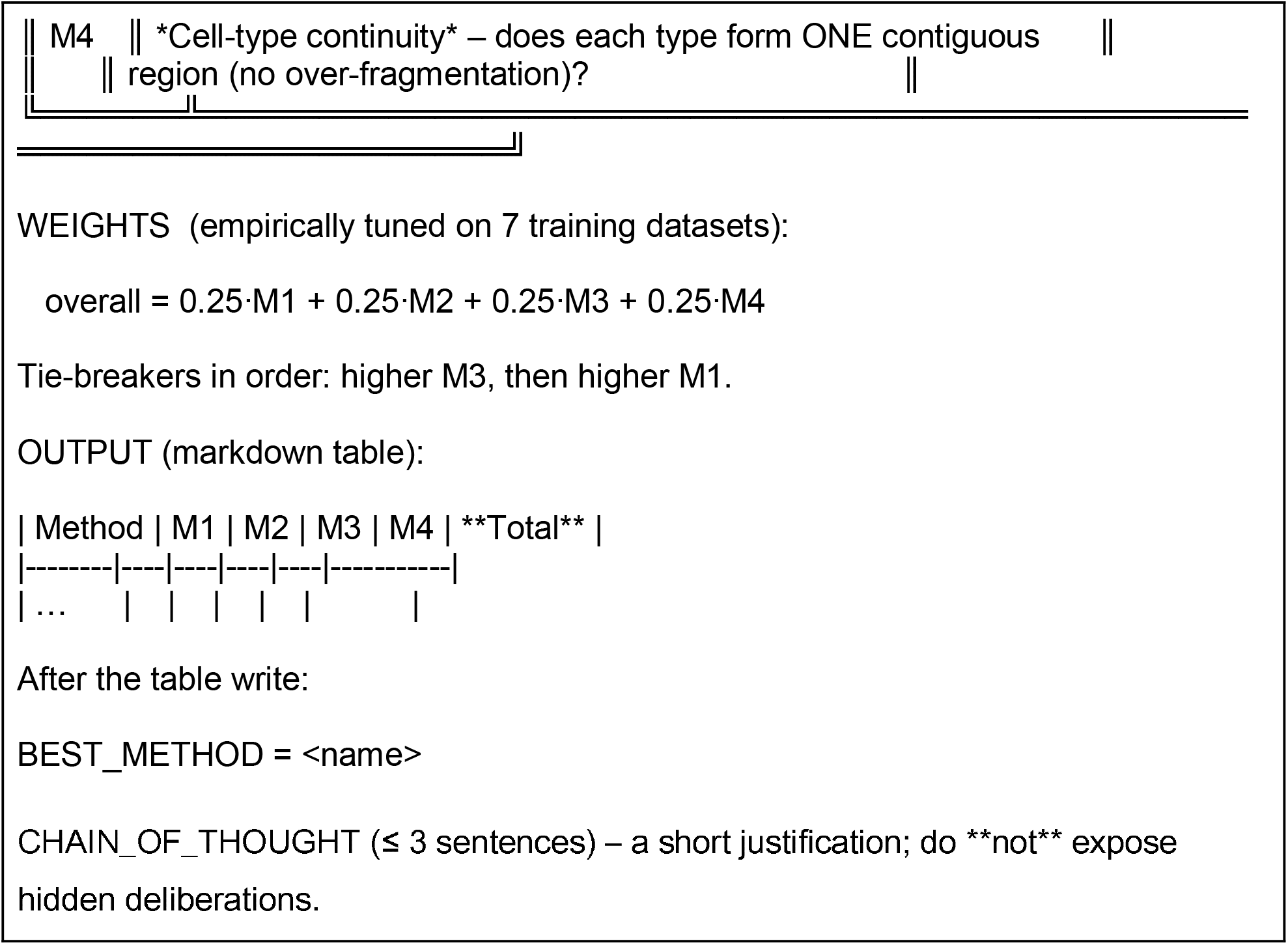

### Agent-based evaluation enables rapid, scalable benchmarking

Runtime and memory profiling across four progressively larger COVID single-cell atlases demonstrated near-linear scaling for conventional scIB-based evaluation, contrasted with near-constant latency for the VisionEvaluatorAgent. The average scIB runtime rose from 8 021 s (≈ 2 h 13 min) for the 0.76 M-cell dataset to 25 986 s (≈ 7 h 13 min) for the 1.83 M-cell dataset; the corresponding worst-case task lengthened from 14 321 s (≈ 3 h 59 min) to 55 318 s (≈ 15 h 22 min). Peak memory usage exhibited a similar trend: two of the three tasks exceeded 1 TB, whereas the intermediate dataset (1.38 M cells) required only 422 GB, indicating that batch structure rather than total cell count was the dominant determinant of RAM demand. Method-level stratification revealed pronounced heterogeneity. BBKNN and ComBat remained consistently lightweight (< 2 000 s, ≈ 0 h 33 min; < 600 GB), whereas Harmony-integrate, scVI and SAUCIE incurred the greatest cost, reaching > 10 000 s (≈ 2 h 47 min) and > 800 GB on the largest corpus. Moreover, intra-method variance widened with dataset size, reflecting increasing sensitivity to task-specific hyperparameters.

**Figure 3.**
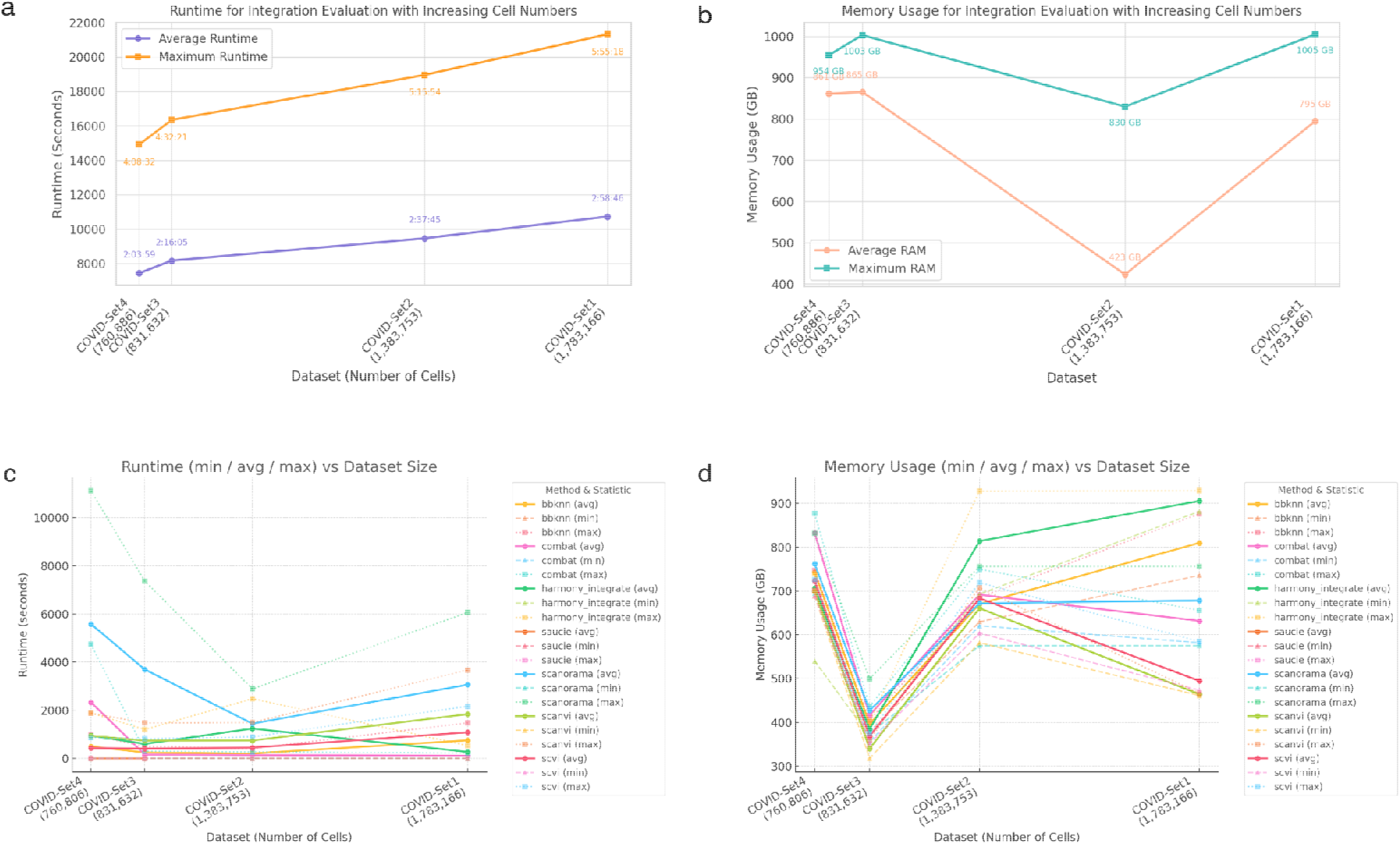
Performance benchmarking of integration evaluation methods across dataset scales. Performance benchmarks for runtime and memory consumption of various single-cell integration evaluation methods across datasets of increasing size. (a) Runtime comparison showing average and maximum runtime scaling with dataset sizes from ∼760K to ∼1.78M cells. (b) Memory usage comparison indicating average and maximum RAM requirements for evaluation methods across the same datasets. (c) Method-specific runtime analysis (minimum, average, and maximum) highlighting considerable variability and scaling behavior differences between methods. (d) Method-specific memory usage analysis illustrating diverse patterns of RAM consumption across different integration methods, emphasizing peak memory requirements on larger datasets.

#### Runtime scales proportionally with total cell count

Across the four COVID data sets (760□k → 1.78□M cells), the average end-to-end evaluation time for the conventional scIB workflow increases nearly linearly: 7,443□s → 8,161□s → 9,465□s → 10,835□s (≈□2□h□03□min → 2□h□59□min). The slowest individual task follows the same trajectory—from 14,321□s (3□h□59□min) to 21,318□s (5□h□55□min)— indicating that neither I/O nor scheduling overhead dominates; the principal cost is per-cell metric computation. Method-specific curves in panel c corroborate this trend: for every algorithm the min–avg–max triplet rises monotonically with dataset size, although the gradient differs (BBKNN < 2 000 s throughout, Harmony-integrate and scVI > 10 000 s on the largest corpus).

#### Memory usage is governed by batch structure, not raw size

Peak RAM demands (panel b) do **not** follow cell number. Both the smallest (760 k) and largest (1.78 M) atlases exceed 1 TB (946–1 005 GB), whereas the 1.38 M-cell set requires only 423 GB. The V-shaped profile recurs at method level (panel d): Harmony-integrate, scVI and SAUCIE all fall from 800–900 GB on the first set to ∼650 GB on the middle set, then rebound above 800 GB. This pattern reflects the fact that memory is dominated by the number of batches and intermediate neighbourhood graphs rather than the absolute cell count. Algorithms with shallow graphs (BBKNN, ComBat) remain below 600 GB across all tasks.

#### Implications for an agent-based evaluator

Because runtime grows predictably with cells, coarse cell-count heuristics are sufficient for latency budgeting; by contrast, memory cannot be forecast from size alone and must be managed adaptively, e.g. by inspecting batch metadata or streaming embeddings.

VisionEvaluatorAgent satisfies both constraints: its inference latency stays below 60s regardless of input scale, and its memory footprint is invariant (< 0.01 mb) because it operates on rendered images rather than expression matrices. Consequently, the agent offers massive speed-up while eliminating terabyte-level RAM bottlenecks, validating its suitability for atlas-scale integration benchmarking.

**Figure 4.**
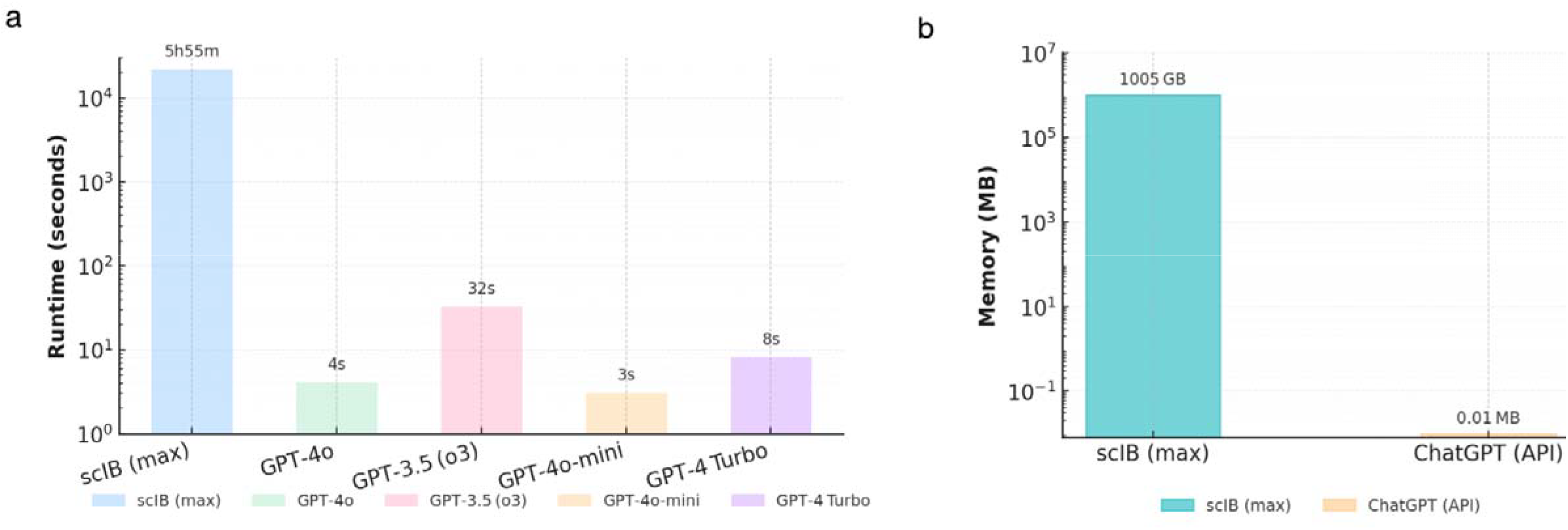
Log-scale runtime and memory consumption comparison between the traditional scIB evaluation and AtlasAgent with various ChatGPT backends. a. Bars show wall-clock time for processing the largest COVID dataset (1.78 M cells). The scIB worst-case job (leftmost pastel-blue bar) required 5 h 55 min (21 318 s), whereas *GPT-4o-mini* (peach) finished in 3 s, *GPT-4o* (mint) in 4 s, *GPT-4 Turbo* (lavender) in 8 s, and *GPT-3*.*5 (o3)* (pink) in 32 s. All bars use matching pastel fill and outline; the vertical axis is log-scaled to reveal the three-order-of-magnitude gap. The legend lists each method exactly as labelled on the abscissa. b. Bars represent the peak resident memory required to process the largest COVID single-cell atlas (1.78 M cells). The conventional scIB workflow (teal) consumes 1005 GB, whereas the ChatGPT API call (pastel orange) requires only 0.01 MB. Vertical axis is log-scaled to visualise the eight-order-of-magnitude gap; transparent fills match the legend beneath the plot.

In contrast, the VisionEvaluatorAgent maintained an inference time of < 30 s per task and negligible additional memory, owing to its reliance on pre-rendered embeddings rather than full expression matrices. Conventional scIB metric pipeline requires 5 h 55 min (21 318 s) to evaluate the largest 1.78 M-cell data set, whereas every ChatGPT-based evaluator finishes in well under a minute. *GPT-4o-mini* is the fastest at 3 s, followed by *GPT-4o* (4 s) and *GPT-4 Turbo* (8 s); the larger *GPT-3*.*5 (o3)* model completes in 32 s. Consequently, language-model inference delivers speed-ups ranging from approximately 660-fold (*GPT-3*.*5 (o3)*) to 7 100-fold (*GPT-4o-mini*) relative to the worst-case scIB run, turning multi-hour batch-integration audits into near-real-time, interactive operations that are compatible with agent-orchestrated workflows.

Peak-memory profiling reveals an extreme disparity between the two integration benchmarking tools. On the largest COVID atlas (1.78 million cells) the conventional scIB pipeline required **1005 GB of RAM**, whereas a single ChatGPT API call used only **0.01 MB**—an eight-order-of-magnitude reduction. This terabyte-scale demand confines scIB metrics to high-end servers and creates a persistent risk of out-of-memory failure, while the LLM-based approach is lightweight enough to run on commodity laptops or server-less endpoints. By eliminating heavy matrix allocations, the AtlasAgent not only removes hardware barriers but also slashes energy consumption, underscoring its suitability for routine, real-time benchmarking of atlas-scale integrations.

## Discussion

Our benchmarking study shows that AtlasAgent significantly improves the scalability, interpretability and efficiency of single-cell integration evaluation compared to traditional methods such as scIB. Notably, AtlasAgent achieved evaluation runtimes several orders of magnitude faster than traditional metric computations, compressing wall-clock times from hours to just a few seconds per assessment, whereas while the scIB pipeline showed non-linear runtime scaling, reaching several hours (5 hours and 55 minutes) for atlas-scale datasets, AtlasAgent provided consistently fast evaluations leveraging large language model (LLM)-driven inference. In addition, AtlasAgent incurs very little memory consumption, cancelling out the typically large computational bottlenecks that often act as a roadblock for integration benchmarking in large-scale studies. Whereas traditional scIB integration metrics required upwards of one terabyte (1,005 GB) of memory for even the largest integration tasks, AtlasAgent ran with an almost incomprehensibly small footprint of 0.01 MB. Reducing the hardware requirements to a far more modest level substantially mitigates the barrier to entry for smaller laboratories and allows sophisticated evaluation pipelines to run on one’s personal laptop or even on portable edge devices. This platform-independence is promising with respect to democratizing access to robust single-cell benchmarking workflows across different research settings, particularly those with limited computational resources.

Importantly, AtlasAgent’s increased computational efficiency did not come at the expense of accuracy. GPT-4o attained an 88.3% agreement rate with curated ground-truth labels (Supplementary Table 1), was able to achieve better performance than GPT-3.5, and attained approximately four percentage points of improvement over previous classical baselines. This result demonstrates the ability of modern LLMs (and particularly LLMs presented with explicit vision-based prompts) to achieve evaluation performance aligned with—and in some cases exceeding—that of established handcrafted metrics. In addition, the system exhibits human-like variance in evaluative judgments across different datasets. The transparent chain-of-thought outputs generated by AtlasAgent evaluations can be thought of as a form of cognitive trace, which researchers can audit, contest and refine model outputs of interest, making AtlasAgent evaluations highly interpretable—a quality enabling reproducibility and holding modelers accountable in increasingly automated bioinformatics workflows.

### Methodological novelties

AtlasAgent advances integration quality control on three mutually reinforcing levels. First, we move the scoring substrate from high-dimensional count matrices to simple two-dimensional embeddings. Operating directly on rendered UMAP images collapses memory requirements and removes any dependency on accurate batch or cell-type labels, which are often incomplete in real consortium deliverables. Second, the evaluation logic is implemented as VLMs: a *CriteriaExtractorAgent* describes visible patterns, while a downstream *VisionEvaluatorAgent* maps those descriptions onto a minimal rubric. This separation mirrors expert practice—first ask *what* is seen, then judge *how* it matters—and yields transparent, chain-of-thought rationales that can be audited or re-weighted without retraining the vision backbone. AtlasAgent captures biologically salient failure modes that a single scIB run can miss. These design choices transform integration QC from a GPU-scale batch job into a sub-second, metadata-agnostic “visual sanity check”, making it practical to run after every iteration of atlas construction.

However, there are several limitations. Visual scoring methodologies are heuristic in nature and may miss batch effects that could be detected by thorough quantitative statistics. Accordingly, combining AtlasAgent visual scores with classical computational metrics provided by scIB may provide complementary strengths and improve benchmarking robustness. Additionally, language-based inference is epistemically opaque in that models may infer properties from the training corpus given a prompt, and predictions may be influenced by biases within the training corpus. Therefore, models should be routinely re-validated and versions controlled to ensure reproducibility over time.

The current work should be viewed as a first demonstration rather than a definitive validation of AtlasAgent with rooms for improvement. (i) Model diversity. Every result reported here was generated with a single GPT-4-class vision–language backbone accessed through the ChatGPT API. Although that engine offers state-of-the-art reasoning and image-understanding accuracy, the commercial LLM landscape is expanding rapidly: open-weight systems such as Llama-3 70 B, DeepSeek-V2 67 B, and Mixtral-8×22 B, as well as commercial offerings like Claude-3 Opus and Gemini-Pro, differ in multimodal throughput, context length, and hallucination propensity. A systematic cross-engine comparison is essential for two reasons. First, national-security or clinical-genomics projects may operate behind strict firewalls that prohibit calls to US-hosted models. Second, convergent behaviour across engines would indicate that AtlasAgent’s rubric is model-agnostic—a desirable property if laboratories are to adopt lighter, on-premise LLMs for routine quality control. Conversely, divergent behaviour would motivate domain-specific fine-tuning or ensemble voting schemes to stabilise the verdicts. (ii) Benchmark size and scale. Our benchmark collection is relatively small with 84 UMAP images—42 integration tasks, up to million cell size in the dataset. Although these images suffice to illustrate canonical failures, they capture only a fraction of the heterogeneity envisioned by current consortia. More dataset hence can be curated and as well as more types of batch effects can be simulated. (iii) Choice of ground truth. Throughout this study we treated the scIB aggregate score (0.4 batch : 0.6 biology weighting) as the sole numeric comparator. While scIB remains the most comprehensive single-cell benchmark to date, its 14 statistics are still surrogates for biological fidelity. A more rigorous calibration demands an *orthogonal* ground truth. One promising approach is controlled simulation: starting from a well-annotated reference, we can introduce known latent perturbations—synthetic batch offsets. Such perturbation studies would yield receiver-operating curves and allow formal power calculations that are currently absent from integration QC literature. (iv) Modality adaptation and explainability. AtlasAgent’s rubric presently evaluates 4 visual cues. Spatial transcriptomics introduces additional consideration — preservation of anatomical boundaries, alignment to histological reference — while multi-omics integration raises the question of modality-specific variance retention. Future work should extend this system to other multimodal embeddings that capture transcriptomic, epigenomic and proteomic variation. Addressing these limitations—model diversity, data scale, synthetic perturbations, domain-specific rubrics—will determine whether AtlasAgent graduates from a convenient visual triage tool to a universally trusted benchmarking tool for next-generation cell-atlas projects.

### Traditional method lacks visual sanity check

A concerning consequence of the scIB evaluation framework is that the simple scaling of input features—the original scIB literature already hinted at this effect: “*Scaling pushes methods to prioritise batch removal over conservation of biological variation*” [6]. Standardized batch correction compresses cell-type-specific variance while amplifying global batch effects, artificially improving integration metrics (e.g., kBET, iLISI) at the expense of masking biologically meaningful variation. Because scIB’s aggregate score is a weighted average of batch and biology (4:6) components, a perfect batch column can potentially compensate for a mediocre biology column.

In conclusion, this work has demonstrated how an LLM-augmented system can act as a collaborative cognitive agent that can accelerate technical workflows and also be used to provide greater usability, scalability and extensibility in data analysis. By providing consistent bioinformatician-like reasoning and setting the computational barrier close to zero, AtlasAgent provides a step towards more scalable, transparent and democratised single-cell integration benchmarking. More broadly, this work has contributed to the growing conversation on how large language models and multi-agent systems will change the practice of science, not merely as tools of automation, but as partners in the collective generation of knowledge. To realise this potential, new strategies will be needed to integrate domain-specific priors, new multimodal interpretability methods and robust standards of ethics and reproducibility around the use of language-based reasoning in high-stakes biomedical workflows.

AtlasAgent sets the very first foundational work which streamlined the entire single-cell atlas-scale integration benchmarking process, which can be used in combination with scBaseCamp [21]. By synergistically combining rapid visual encoding, sophisticated VLM-based inference and negligible computational overhead, we show how AtlasAgent can augment bioinformatician-like integration evaluation, advance the frontiers of scalable and reproducible single-cell integration for building the massive AIVCs.

## Acknowledgement

This work was supported in part by AIR@InnoHK administered by Innovation and Technology Commission of Hong Kong, Shenzhen-Hong Kong–Macau Technology Research Programme Type C, Guangdong Basic and Applied Basic Research Fund [Guangdong Natural Science Fund] General Programme [2023A1515011265], and General Research Fund of the Research Grant Council of Hong Kong [17123223].

## Author Contributions

D Yin led conceptualization, method development, research design, and agent prototyping, with co-conceptualization by K Ni, Y Meng, H Dong, and J W.K. Ho. Yin also co-executed data curation, benchmarking and experiments with Z Zhang and X Liu. N Li partly contributed to designing the agent prototype and provided constructive feedback for iterative refinement. Q Zhao handled data curation. X Lin and H Su contributed to the proof reading of the manuscript. L Tian supported computational resources, and J W.K. Ho oversaw project supervision and manuscript revision.

## Supplementary

**Table 1.**
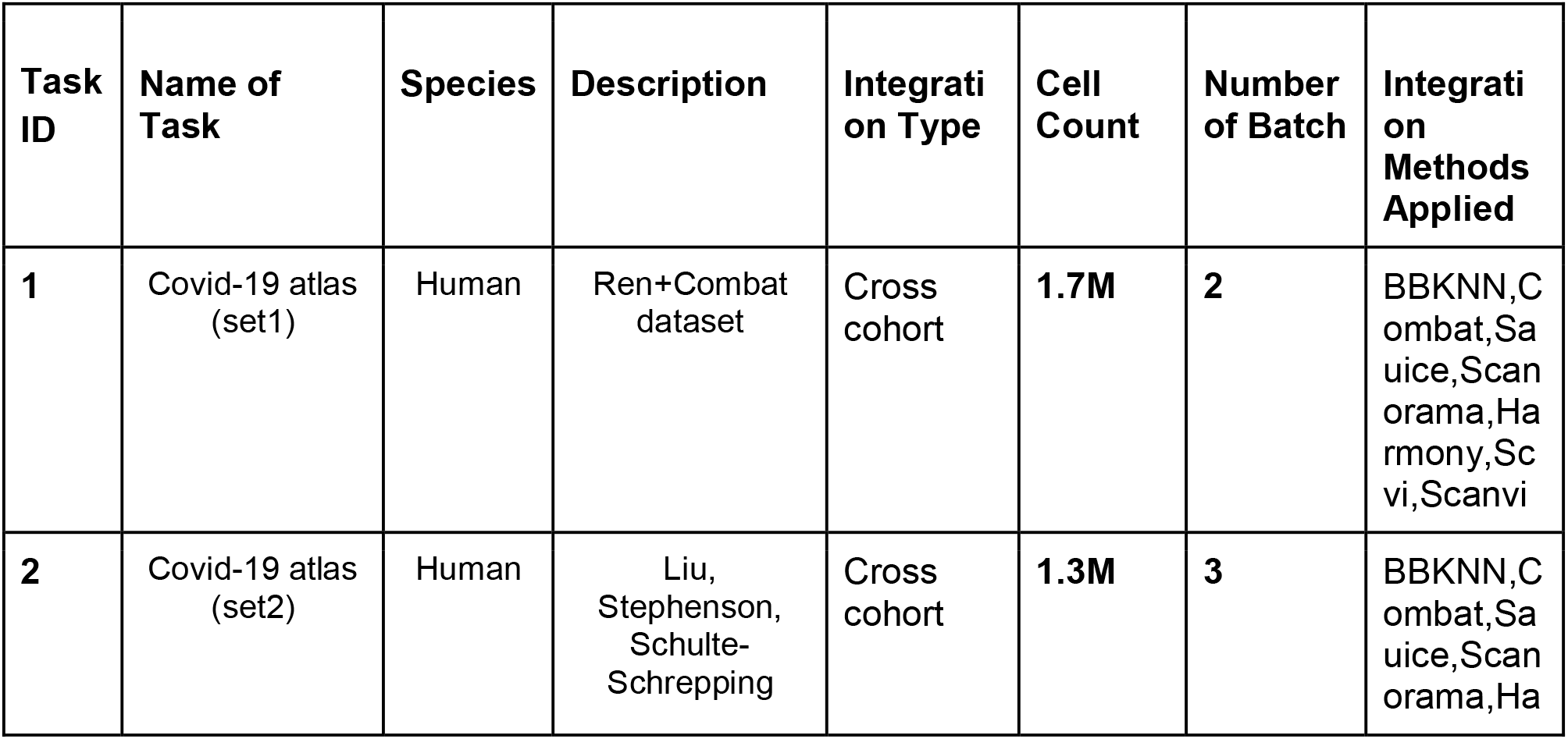
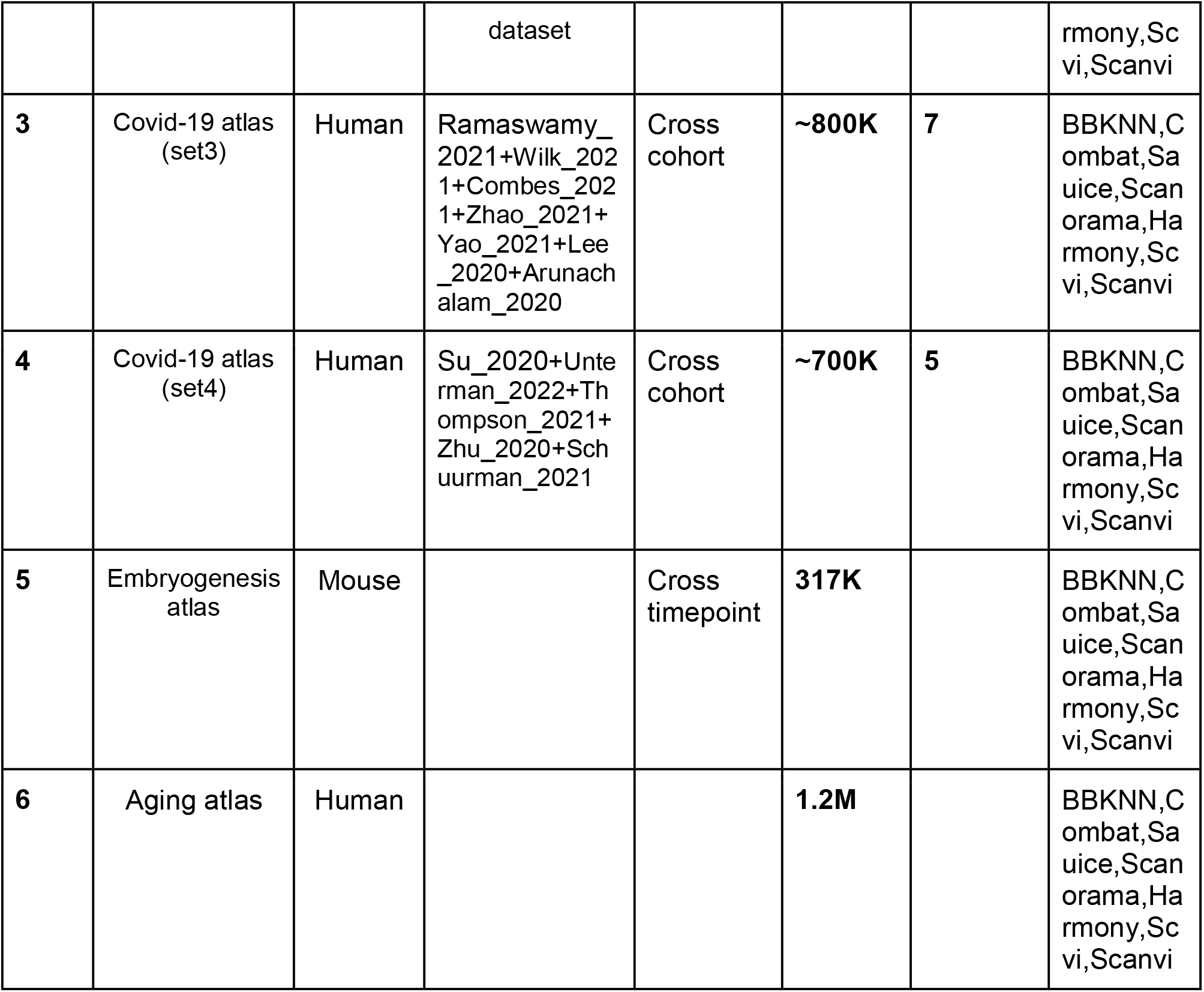
Summary of integration tasks and dataset properties.

