## Supplementary Data 1 for "AtlasAgent: Vision language model and Agent-guided Framework for Evaluation of Atlas-scale Single-cell Integration"

### Acknowledgement

Conceptualization: D Yin, N Ke, M Ye, D Yin, J WK Ho; Method development D Yin, Xi Liu; Research design D Yin,

### Supplementary

Table 1. Summary of integration tasks and dataset properties.

| Task ID | Name of Task | Species | Description | Integration Type | Cell Count | Number of Batch | Integration Methods Applied |
|---|---|---|---|---|---|---|---|
| **1** | Covid-19 atlas (set1) | Human | Ren+Combat dataset | Cross cohort | **1.7M** | **2** | BBKNN,Combat,Sauice,Scanorama,Harmony,Scvi,Scanvi |
| **2** | Covid-19 atlas (set2) | Human | Liu+Stephenson, Schulte-Schrepping dataset | Cross cohort | **1.3M** | **3** | BBKNN,Combat,Sauice,Scanorama,Harmony,Scvi,Scanvi |
| **3** | Covid-19 atlas (set3) | Human | Ramaswamy_2021+Wilk_2021+Combes_2021+Zhao_2021+Yao_2021+Lee_2020+Arunachalam_2020 | Cross cohort | **~800K** | **7** | BBKNN,Combat,Sauice,Scanorama,Harmony,Scvi,Scanvi |

| | | | | | | | |
|---|---|---|---|---|---|---|---|
| **4** | Covid-19 atlas (set4) | Human | Su_2020+Unterman_2022+Thompson_2021+Zhu_2020+Schuurman_2021 | Cross cohort | **~700K** | **5** | BBKNN,Combat,Sauice,Scanorama,Harmony,Scvi,Scanvi |
| **5** | Embryogenesis atlas | Mouse | | Cross timepoint | **317K** | | BBKNN,Combat,Sauice,Scanorama,Harmony,Scvi,Scanvi |
| **6** | Aging atlas | Human | | | **1.2M** | | BBKNN,Combat,Sauice,Scanorama,Harmony,Scvi,Scanvi |
